# Belowground plant-plant signaling of root infection by nematodes

**DOI:** 10.1101/2020.08.19.254987

**Authors:** Peihua Zhang, Dries Bonte, Gerlinde B. De Deyn, Martijn L. Vandegehuchte

## Abstract

Communication between plants mediated by herbivore-induced volatile organic compounds has been extensively studied aboveground. However, the role of root herbivory in belowground plant-plant communication is much less understood. We here investigated whether root herbivores can trigger plant roots to emit warning signals to neighbouring plants that are not yet in direct contact with them.

We used a split-root system and infected half of the roots of *Agrostis stolonifera* plants with root-knot nematodes (*Meloidogyne minor*) and left the other half uninfected. As a control, we grew plants without nematodes in separate pots. Leachates from each split-root soil compartment and from soils with control plants were applied to separate pots with *A. stolonifera* plants, of which biomass allocation and morphological traits were measured one month after leachate addition.

Plants receiving leachates from the soil with the nematode-free roots of the nematode-infected plants showed a significantly larger total biomass, more root branches, and deeper rooting than plants receiving leachates from the soil with the nematode-infected roots or from soil with control plants. Plants were taller and the root/shoot ratio was higher in plants receiving leachates from soil with the nematode-free roots than in plants receiving leachates from soil with nematode-infected roots. Shoot tiller number was higher in plants receiving leachates from either soil of the nematode-infected plants than in plants receiving control leachates.

Our results suggest that an overcompensation response was triggered by systemically induced root-derived compounds from nematode-free roots of a plant locally infected with root-feeding nematodes. Signals from directly attacked roots of the same nematode-infected plant only caused receiver plants to develop more shoot tillers, possibly for future stolon development to grow away from the infected area. This may indicate an anticipatory tolerance response to root feeders that are still distant and an additional generalized escape response to root feeding.

## Introduction

Plants, as primary producers and sessile organisms, are constantly under attack by a variety of natural enemies. When plants are exposed to herbivory or infection, both constitutive and induced defenses can protect the attacked organs. These defensive strategies include direct resistance via chemical or mechanical traits that reduce herbivory, indirect resistance via traits enhancing the action of enemies of the attacking herbivores, or tolerance such as regrowth after damage (Rasmann and Agrawal, 2008).

A defence that is induced in response to local attack is often established systemically throughout the whole plant (Heil and Ton, 2008; Mithöfer and Boland, 2012; Ross, 1961). Systematically induced resistance can be achieved by two mechanisms: systemic transport of defensive metabolites or *de novo* expression of defensive metabolites that is activated by translocated signals from the stress-exposed tissues (reviewed by Heil and Ton, 2008). Either way, these induced defensive compounds can be found in distal organs which have not yet been damaged by herbivores. Furthermore, plants can shift biomass and resources away from attacked organs as a compensatory adjustment to cope with herbivory both above- and belowground (Babst et al., 2005; Gómez et al., 2012; Hanik et al., 2010; Henkes et al., 2008; Newingham et al., 2007).

Communication between plants has been extensively studied above ground, with a focus on HI-VOCs (herbivore-induced volatile organic compounds; Karban *et al.*, 2014; Karban, Yang and Edwards, 2014). These compounds are released as plants are wounded by herbivores and can be directly repellent to herbivores or attractive to predators and parasites of herbivores that can decrease levels of damage inflicted by herbivory (reviewed by Karban, Yang and Edwards, 2014). In addition, VOCs can be perceived by proximate neighbours that eavesdrop on the volatile signals emitted by the herbivore-damaged plant. In response to the “signal” in the volatile, receiver plants can start expressing genes and synthesizing defensive proteins and phytohormones involved in plant defenses (reviewed by Delory *et al.*, 2016) or can prime their defenses against pests (Frost *et al.*, 2008; Heil and Kost, 2006).

Upon herbivore attack, root exudates can also mediate positive plant-plant interactions by production of secondary metabolites to increase herbivore resistance in exudate-exposed neighbouring plants or by reducing the herbivore populations by attracting predators and parasites of the attacking herbivore (reviewed by Bais *et al.*, 2006; Dicke and Dijkman, 2001; Guerrieri *et al.*, 2002). Such root exudates also play significant roles in stomatal aperture by sending water stress cues (Falik *et al.*, 2012) and in kin recognition by carrying information about the degree of genetic relatedness (Semchenko et al., 2014). However, the effects of root herbivory on plant-plant interactions mediated by root exudates have been studied less than the role of leaf VOCs in conferring resistance against aboveground herbivores.

Root-knot nematodes of the genus *Meloidgyne* are obligate plant parasites that must enter into the roots to feed and reproduce (Williamson and Gleason, 2003). They reduce plant growth by formation of galls and giant feeding cells in the roots which can result in a deformed root system and substantial plant growth suppression (Abad et al., 2003). Studies have shown that root feeding by endoparasitic nematodes can induce systemic changes in gene expression and defense responses across plant compartments of both dicots and monocots (reviewed by Biere and Goverse, 2016). Root infection by root-knot nematodes can increase the level of defence compounds in the root exudates (Wurst et al., 2010). These systematically induced defensive compounds can also be found in distant undamaged plant parts (Mithöfer and Boland, 2012). However, our understanding of defensive responses of plants against plant-parasitic-nematodes is mainly based on molecular studies focusing on the organismal level. In contrast, few studies have focused on the ecological consequences of plant-nematode interactions for neighbouring plants (Liu and Park, 2018). Thus, whether root exudates from a plant infected with root-knot nematodes can induce a plastic response in neighbouring plants remains an open question. Therefore, we used a model system to test our hypothesis that root-feeding nematodes trigger a systemic response such that root exudates from non-damaged roots of the infected plant trigger plant growth responses in neighboring non-infected plants. We more specifically hypothesized that: 1) Leachates from soil containing nematode-infected and nematode-free roots of the same nematode-infected individual plant would cause a similar response of the receiver plants due to the induced systemic defense in the entire plant; and 2) Plants receiving leachates from either nematode-infected or nematode-free roots of the same plant would allocate less biomass belowground than plants receiving leachates from uninfected control plants to avoid root damage from future nematode infection, similar to the response of nematode-infected plants that allocate resources away from attacked organs.

## Material and methods

### Donor Plants

Creeping bentgrass seeds were obtained from a commercial supplier (Cruydt-Hoeck, Netherlands). Seeds were first surface sterilized. They were treated with 3% household bleach for 10 minutes, rinsed ten times with distilled water, treated with 10% ethanol for another 10 minutes and rinsed another ten times with distilled water (Vandegehuchte et al., 2010). The surface-sterilized seeds were placed on wet filter papers in a Petri dish (50 seeds/dish) and left to germinate at 22°C in a plant breeding room with a light/dark regime of 16 h/8 h. Two-week-old seedlings with an average shoot length of 1.5 cm and average root length of 4 cm were transplanted into 500 ml pots containing sand (Decor Son, Netherlands) to grow for another seven weeks until reaching a satisfying root system for root-splitting.

### Split-root system

Two 10.8×8.2×9.3 cm^3^ (length×width×height) pots were taped together to make the split-root system. There were 12 holes at the bottom of the pots for later leachate collection. A layer of cotton (10.5×8cm^2^) were placed at the bottom of each pot to prevent sand leaching and prevent roots from growing out of the pots. Sand (Decor Son, Netherlands) was sterilized by autoclaving at 121°C for 30 minutes before adding into the pots. The root system of nine-week-old *A. stolonifera* plants were carefully divided into two halves as similar as possible and transplanted into the split-root system. The pot with the half of the roots to be inoculated with root-knot nematodes was labeled “N (+)”. The pot with the other half of the root system, which was not to be inoculated with nematodes, was labeled “N (-)”. We installed 30 such split-root systems in total. Separate seedlings of the same age without nematodes or root splitting were transplanted into 10.8×8.2×9.3 cm^3^ (length×width×height) pots as control plants (Fig. 1). Plants in split-root systems and control plants were left to grow for another 4 weeks before leachate collection.

**Figure 1:**
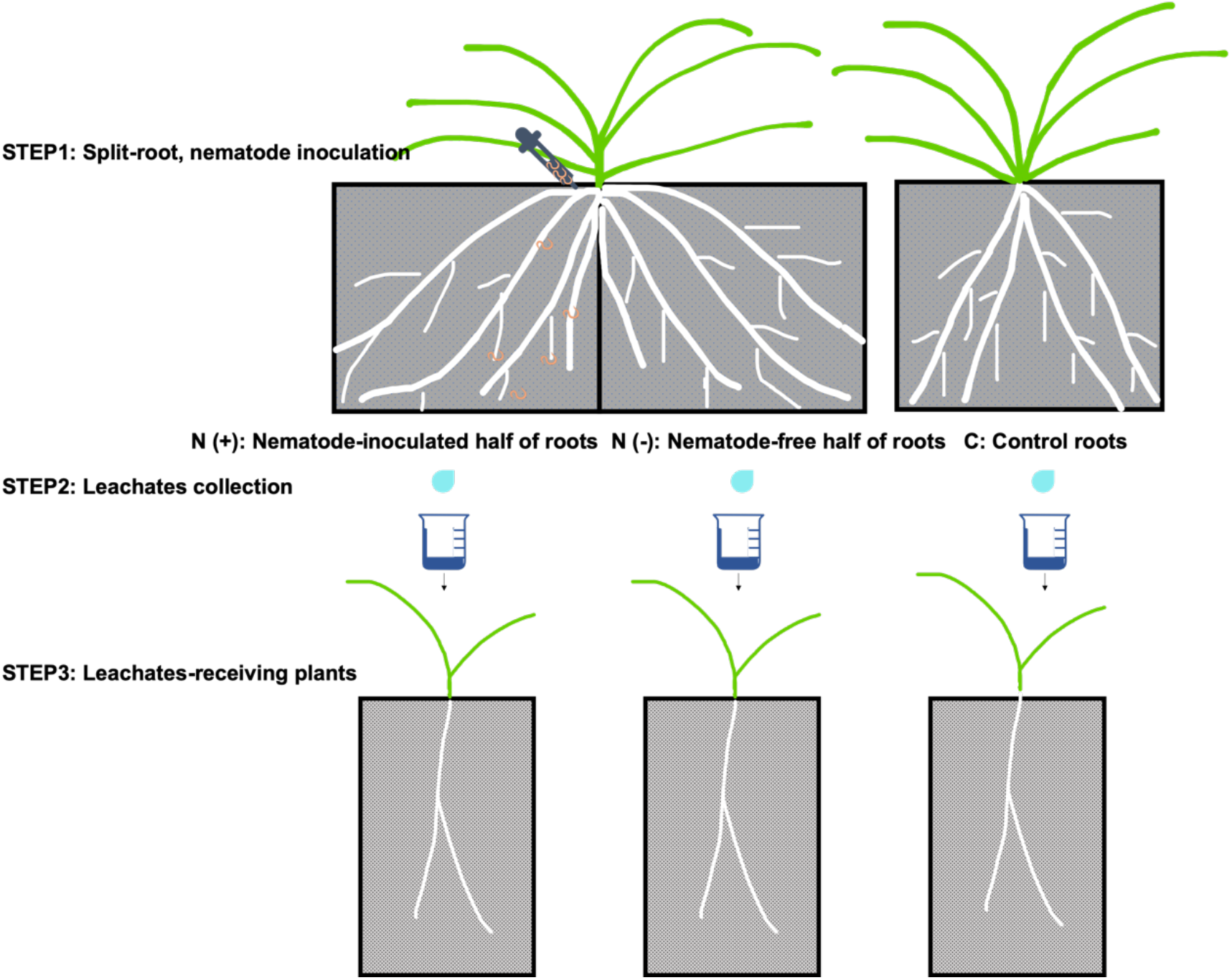
Schematic diagram of the experiment: Step 1: Each plant in a split-root system was inoculated with root-feeding nematodes in one compartment containing half of its roots, while the other compartment containing the other half of the roots was left nematode-free. Nematode-free plants served as controls. The three treatment levels were thus: nematode-infected roots, nematode-free roots from the same nematode-infected pant, and nematode-free roots from a nematode-free control plant. Each pot had the same volume. Step 2: leachates were collected as leachates which were filtrates of water that has percolated through soil with roots. Step 3: leachates were applied to separate receiver plants. Each treatment group has 30 replicates for leachate collection and 30 receiver plants for leachate application, for a total of 90 receiver plants. The response variables measured in the receiver plants were total biomass, root/shoot ratio, shoot length, shoot tiller number, rooting depth and root number.

### Nematode inoculation

Roots in the “N” pots were inoculated with nematodes one week after transplanting the plant into the split-root system and 5 ml of nutrient supplement (2 g of the product in 1L distilled water, COMPO, NPK:16-9-20) was added to each pot in each split-root system and each control pot before the nematode inoculation. The root-knot nematodes (second-stage juveniles J2) cultured on potatoes were purchased from HZPC, Netherlands. Four ml of the inoculum with a concentration of about 350 J2 individuals/ml was added into a hole at 0.5 cm distance from the stem and 2 cm deep below the soil surface of the treatment plants, i.e. approximately 1400 juveniles per treatment plant. The same amount of water (4 ml) was added into the rhizosphere of the roots in the “R” compartment of the split-root systems and of the control plants.

### Receiver plants

Seeds of *A. stolonifera* from the same batch were surface sterilized as described above and germinated three weeks before leachate collection from the donor plants. Two-week-old seedlings were planted into a 1 L pot (13.5 cm in height) containing the same sterilized sand as the pots with the donor plants. Transplanted receiver plants received 5 ml nutrient supplement (2 g of the product in 1L distilled water, COMPO, NPK:16-9-20). The receiver plants were placed in random positions in another plant breeding room with a light/dark regime of 16 h/8 h and 22°C constant temperature. Leachates were collected and applied to the receiver plants one week after the transplantation. The leachates from 30 control plants, 30 nematode-containing split-root compartments, and 30 nematode-free split-root compartments were collected multiple times and each applied to a single receiver plants, for a total of 90 receiver plants. After one month of repeated leachate application (see next paragraph), the receiver plants were harvested.

### Leachates

Donor plants (split-root plants and control plants) were not watered for three days before each leachate collection. Each week, leachates of the root system were gathered by addition of 120ml of distilled water to the pot (Fenwick, 1949) and collecting 100 ml of the percolated water. Twenty ml of nutrient supplement (2 g of the product in 1L distilled water, COMPO, NPK:16-9-20) were added to each donor plant (10ml per split-root compartment and 20ml per control plant) after the leachate collection and one week before the next collection. The leachates were then filtered through filter paper with a pore size of 20 μm (Whatman, Quantitative filter papers, ashless grades, grade 41) before use to remove nematodes potentially present in the leachates. Filtrated leachates were checked for nematodes under the microscope during the experiment by randomly selecting 15 out of the 30 leachate samples from both N (+) and N (-) roots during the first and last week of leachate collection. No nematodes were present in any of the checked samples. These leachates were then added to the sand of receiver plants. Receiver plants were treated with the filtrated leachates once a week for a total of four weeks. Leachates were applied carefully with a pipette at a distance of 0.5 cm from the stem of the receiver plants. Receiver plants received three quantities (40ml, 40ml and 20ml) of root leachates during the first two weeks, two quantities of 50ml in the third week and one quantity of 100ml in the last week. During the experiment the stocks of leachates were stored in sealed plastic bottles at 4°C for maximum one week.

### Receiver plant traits

Upon harvest, total biomass, root/shoot ratio, shoot height, shoot tiller number, maximum rooting depth and root number were quantified for all receiver plants. Fresh roots were cleaned and scanned using an EPSON scanner (Epson Expression 11000 XL). Image J was used to count the number of first-order roots and measure the maximum root depth by analyzing the scans of the roots. After scanning, root samples were dried at 70 °C for 72 hours before weighing. Fresh shoots were cleaned before measuring the length and number of tillers. Shoot fresh biomass was measured before shoots were dried at 70 °C for 72 hours.

### Statistical analysis

The effects of leachate origin on each trait of the receiver plants were analyzed with linear models, using the lme4 package in R. Data of the shoot length were transformed using Tukey’s Ladder of Powers to produce more normally distributed residuals (transformTukey function, rcompanion package). The function simply loops through lambda values and then chooses the lambda that maximizes the Shapiro-Wilks W statistic. Only the data of shoot length has been transformed with the lambda value of 0.45. All predictors were analyzed as categorical variables. Leachate origin has 3 levels: nematode-inoculated split-root compartment (N), nematode-free split-root compartment (R) and control plants (C). Leachate origin was included as fixed effect and donor-plant identity was included as random effect in our model. Shapiro-Wilk normality tests were used to check for normal distribution of the residuals. Residuals were plotted after analysis and found to be (approximately) normally distributed and homogeneous. Tukey multiple comparison method were used to indicate significant differences among the three treatment groups using package emmeans in R.

## Results

Receiver plants showed significant responses to the leachates from the different split-root compartments and control plants (Table 1). Receiver plants treated with leachates from nematode-free split-root compartments (N (-) plants) produced more biomass in total and both in roots and shoots than plants treated with leachates from nematode-inoculated split-root compartments (N (+) plants), or with leachates from control plants (C plants, Fig. 2a; Fig. 3). The root/shoot ratio of N (-) plants was significantly higher (p=0.002) than that of N (+) plants, while C plants had intermediate root/shoot ratios not significantly different from that of N (+) or N (-) plants (Fig. 2b). Shoots of N (-) plants were significantly longer (p=0.006) than those of N (+) plants, with the latter having the shortest shoots of all leachate-receiving plant groups. The shoot length of C plants was not significantly different from that of N (+) plants nor N (-) plants (Fig. 2c). Both N (+) and N (-) plants had significantly more (p<0.0001; p=0.005) shoot tillers than the C plants, but the number of shoot tillers was not significantly different between N (+) and N (-) plants (Fig. 2d). N (-) plants had significantly deeper (p<0.0001) roots than both N (+) and C plants. The rooting depth of the latter two was not significantly different (Fig. 2e). N (-) plants had significantly more root branches (p<0.0001) than the N (+) and C plants, which had similar numbers of root branches (Fig. 2f).

**Figure 2:**
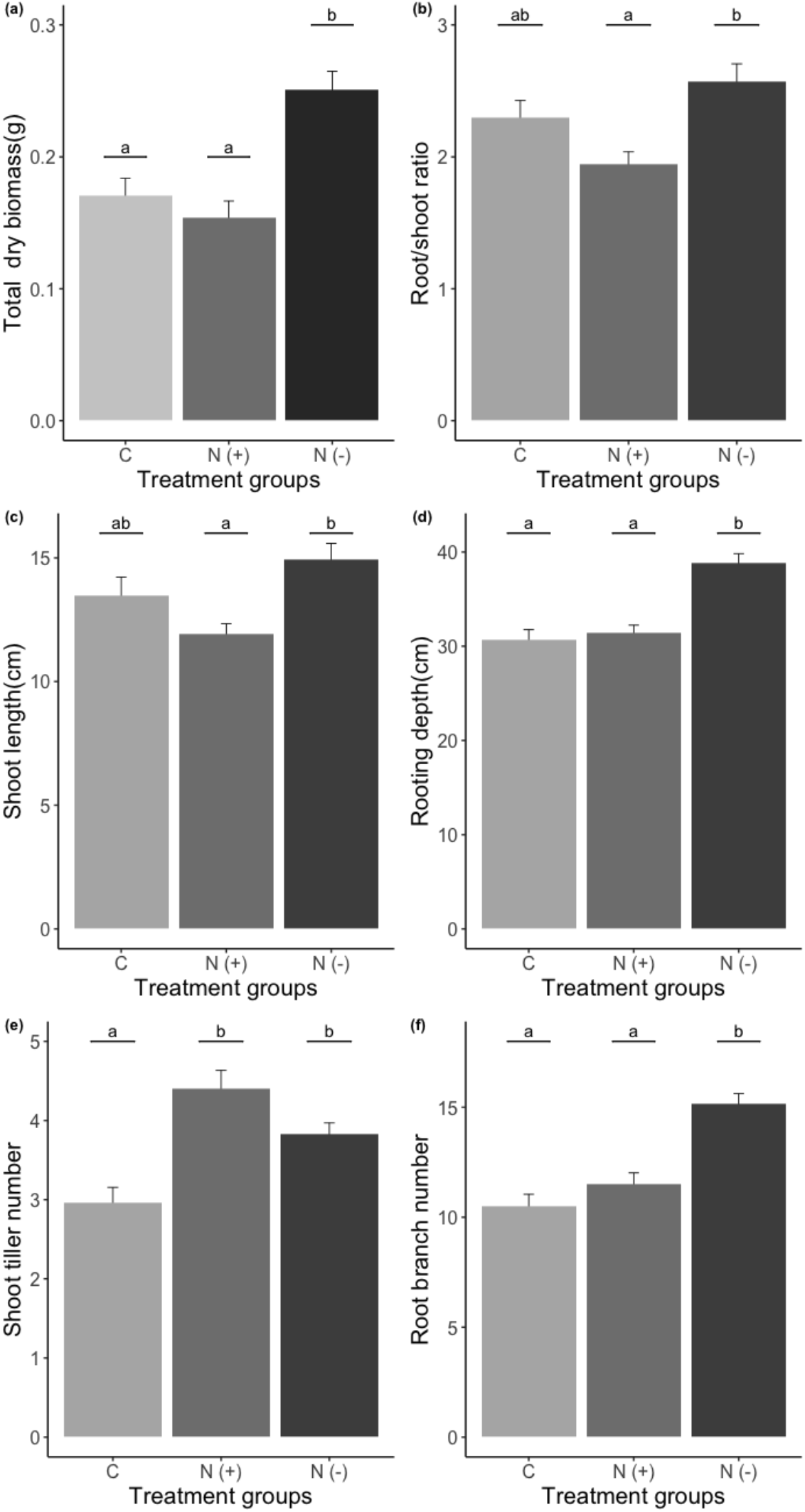
Effect of leachates from different origins (mean ± SE, n = 30) on total dry biomass of the entire plant (a), root/shoot ratio (b), shoot length (c), rooting depth (d), shoot tiller number (e) and root branch number (f) in receiver plants. C indicates receiver plants treated with leachates from control roots with no nematode inoculation; N (+) indicates receiver plants treated with leachates from soils with nematode-infected roots of the nematode-inoculated plants; N (-) indicates receiver plants treated with leachates from soils with nematode-free roots of the nematode-inoculated plants. Different letters above the means indicate significant differences between groups based on Tukey multiple comparison test.

**Figure 3:**
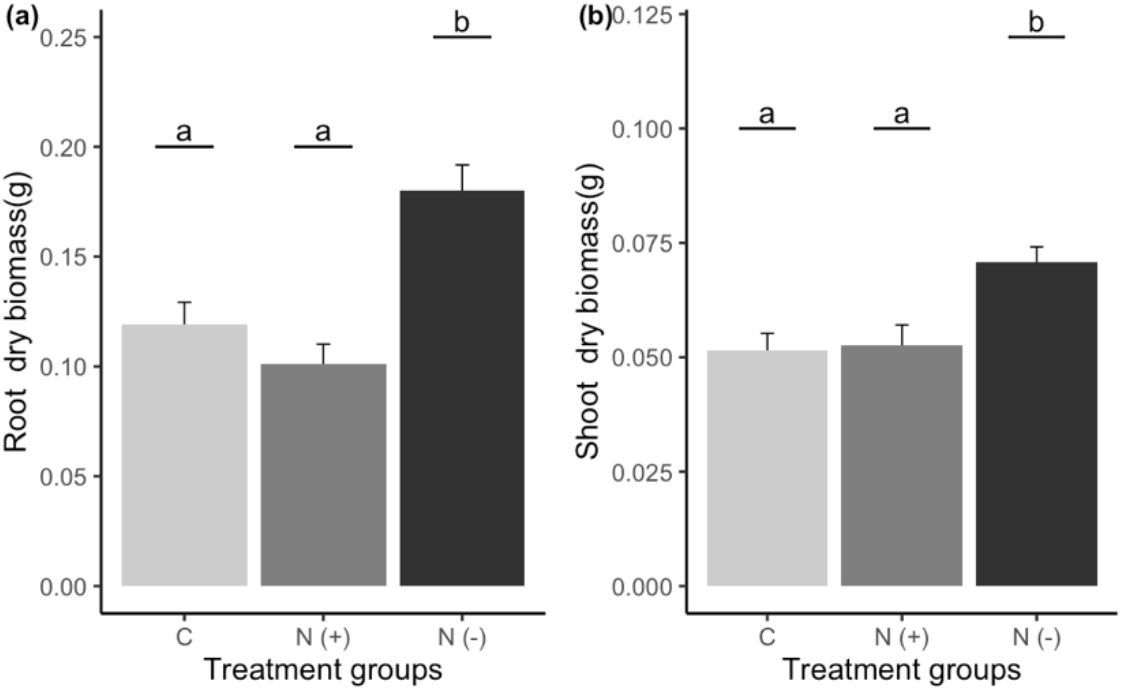
Effect of leachates from different origins (mean ± SE, n = 30) on root dry biomass (a) and shoot dry biomass(b) in receiver plants. C indicates receiver plants treated with leachates from control roots with no nematode inoculation; N (+) indicates receiver plants treated with leachates from soils with nematode-infected roots of the nematode-inoculated plants; N (-) indicates receiver plants treated with leachates from soils with nematode-free roots of the nematode-inoculated plants. Different letters above the means indicate significant differences between groups based on Tukey multiple comparison test.

**Table 1:** Analysis of variance of the effect of leachate treatment of donor plants (control, nematode-infected roots, or nematode-free roots of nematode-infected plant) on measured traits of leachate-receiving plants

| Factor | Num Df | Den Df | F | P |
| --- | --- | --- | --- | --- |
| **<u>Root/shoot ratio</u>** |  |  |  |  |
| Leachate treatment | 2 | 87 | 6.885 | **0.0016** |
| **<u>Root biomass</u>** |  |  |  |  |
| Leachate treatment | 2 | 48.74 | 20.418 | **3.62e-07** |
| **<u>Shoot biomass</u>** |  |  |  |  |
| Leachate treatment | 2 | 50.937 | 8.7515 | **0.00054** |
| **<u>Total biomass</u>** |  |  |  |  |
| Leachate treatment | 2 | 47.514 | 19.253 | **7.51e-07** |
| **<u>Shoot length</u>** |  |  |  |  |
| Leachate treatment | 2 | 87 | 5.378 | **0.0062** |
| **<u>Maximum rooting depth</u>** |  |  |  |  |
| Leachate treatment | 2 | 38.419 | 34.58 | **2.57e-09** |
| **<u>Shoot tiller number</u>** |  |  |  |  |
| Leachate treatment | 2 | 87 | 14.42 | **3.91e-06** |
| **<u>Root number</u>** |  |  |  |  |
| Leachate treatment | 2 | 53.575 | 24.168 | **3.31e-08** |

## Discussion

We used a split-root system to investigate whether leachates from two halves of a root system belonging to the same plant but with different nematode-infection status would induce a similar response in plants treated with these leachates. Contrary to our hypotheses, we show that leachates from the same nematode-infected individual plant have different effect on receiver plants at different scales. At a local scale, plants receiving leachates from soil with nematode-infected roots were significantly smaller in size and allocated relatively less biomass to roots than plants receiving leachates from nematode-free roots of the same infected plant. This difference in plant size and resource allocation pattern indicates a different composition of both leachate types. Nevertheless, plants receiving either type of leachate from nematode-infected plants had a larger number of shoot tillers than plants receiving leachates from uninfected control plants. This suggests that only the number of shoot tillers responded to a generalized, warning signal expressed by the entire root system of a nematode-infected plant. Plants thus showed phenotypically plastic responses to the signal from nematode-free or nematode-infected roots of the same nematode-infected neighbour plant. When a plant grows closely to locally nematode-infected roots from its neighbour plant, it will develop more shoot tillers as a preparation for future very-likely root damage by vicinal nematodes. When a plant has a nematode-infected neighbour plant but is in contact with its not-yet damaged healthy roots, it will increase in size with a large, deep root system and tall and numerous shoots (Fig. 4). Our results indicate that plants can distinguish –and respond to– reliable cues from their soil environment.

**Figure 4:**
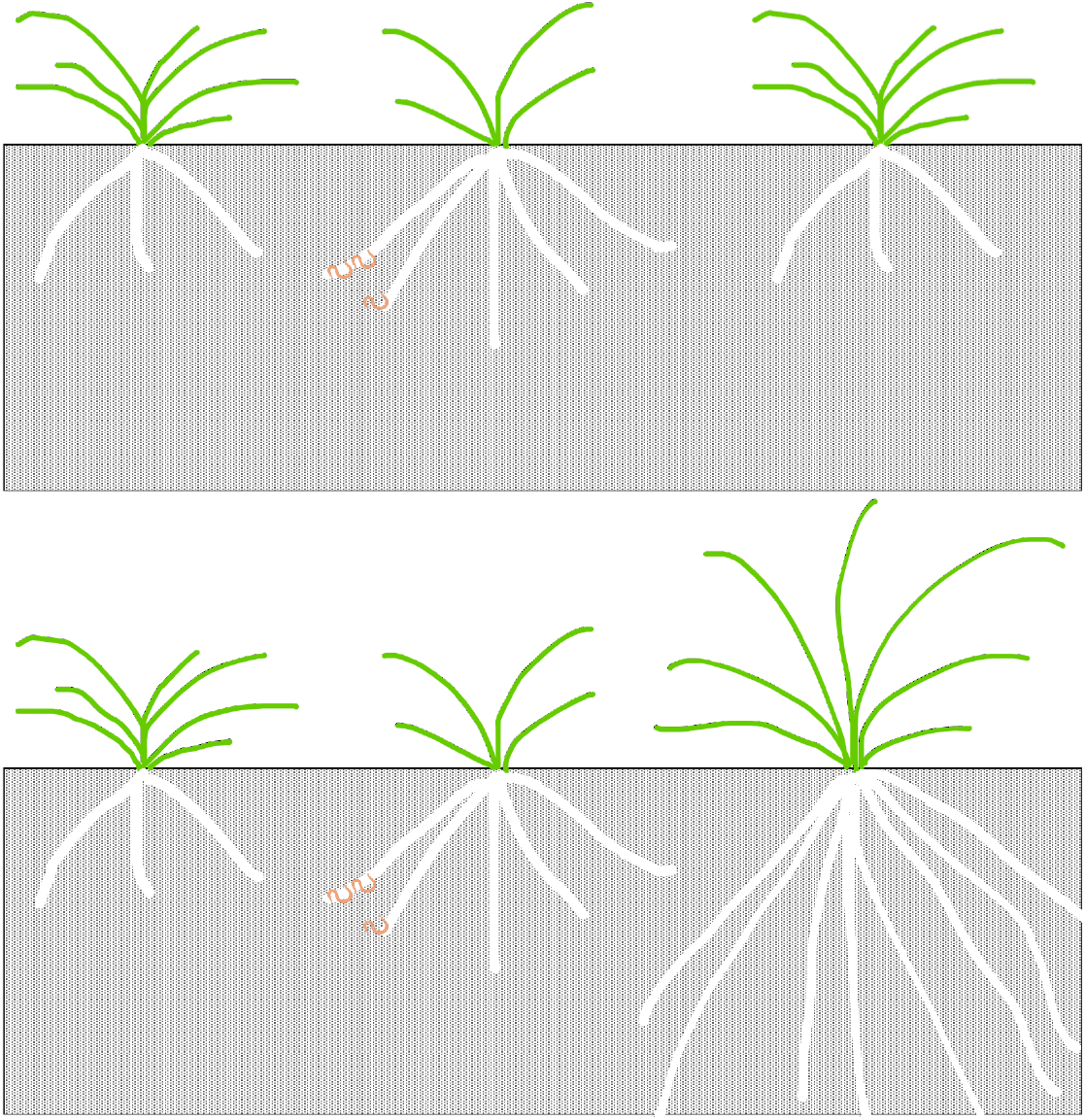
An illustration of the hypothesized plant responses to a nematode-infected neighbour (above): we expected a plant to send a similar warning signal to its neighbouring plants due to the systematic defensive response of the attacked plant; Experimental results (below) showed a different response in the neighbouring plants to a nematode-infected plant: neighbour plants close to the nematode-infected roots tends to grow smaller and those close to the roots with no nematode infection of the same nematode-infected plant tends to allocate more biomass to roots, possibly due to the compositional differences in compounds released from roots with different nematode loads. However, an increased shoot tiller number was a similar response of both types of neighbour plants, regardless of their position with respect to the nematode-infected roots.

It is known that a systemic defense response can be induced in plants after herbivore attack (Heil and Ton, 2008; Mithöfer and Boland, 2012; Ross, 1961). The same defensive compounds can be expected both in the locally attacked site and the distal not-yet-damaged sites. Root infection by root-knot nematodes can cause a different blend of root exudates due to the leaching of ions and metabolic change in the roots (Van Gundy, Kirkpatrick and Golden, 1977). Studies have shown that roots infected with *Meloidogyne* species act as metabolite sinks, which results in increased leakage into the rhizosphere compared with healthy roots (Bird and Loveys, 1975; Dorhout et al., 1993; Haase et al., 2007; Van Gundy et al., 1977). Carbohydrates were the major organic compounds in early exudates and nitrogenous compounds became the major organic compounds after 14 days of nematode infection (Van Gundy et al., 1977). The author also suggested that increased amylase, cellulase and pectinase production by the nematodes before the permanent feeding site was established may contribute to the high level of carbohydrates in the early exudates. Nitrogenous waste products and secreted stylet exudates from the adult female nematodes may contribute to the increase in nitrogenous compounds in the later root exudates. Analysis of stylet exudates from adult female *Meloidogyne incognita* revealed presence of a mixture of proteins in the stylet exudates (Veech *et al.*, 1987). These compounds produced and secreted by root-feeding nematodes play key roles in establishment of feeding sites and activation of defensive responses in plants (Choi and Klessig, 2016; Holbein et al., 2016; Williamson and Gleason, 2003). In our case, leachates from the compartment with nematode-infected roots likely contained nematode-derived compounds. These compounds would then have been taken up by the receiver plant roots, acting as a signal of direct herbivore attack and inducing resource reallocation as in a nematode-infected plant. Studies have shown that direct root herbivory can cause the diversion of resources away from the attacked tissues to organs that are inaccessible to foraging herbivores, such as shoots or stems (Newingham et al., 2007; Robert et al., 2014). On the other hand, leachates from the compartment with nematode-free roots would have lacked these compounds leaking from feeding sites, resulting in a different response of the receiver plants.

Strategies by which plants can defend themselves against herbivory include chemical defenses and regrowth (Ramula *et al.*, 2019). This regrowth, under certain favorable conditions, may produce more biomass in herbivore-attacked plants than in herbivore-free plants, which is also known as overcompensation (Belsky, 1986). Overcompensation can be viewed as an expression of plant tolerance that may be selected for when herbivore damage is predictable and extensive (reviewed by Ramula *et al.*, 2019). However, this overcompensation is often compared quantified as the fitness difference between plants directly damaged by herbivores and plants that are undamaged. Tolerance is the ability of plants to maintain fitness in the presence of stress (reviewed by Heil, 2010). It is also one of plants’ plastic responses to encounters with their enemies. Plants have evolved phenotypically plastics defenses to be prepared for future damage from herbivore stress that is difficult to anticipate. Thus, reliable cues from surrounding environment such as egg deposition and leaf volatiles emitted from attacked neighbors (Doss et al., 2000; Heil and Karban, 2010; Hilker and Meiners, 2006) can be utilized by plants to indirectly increase their resistance to pre-empt upcoming attacks. In our case, plants grew significantly larger after treating them with leachates from soils containing unattacked roots of a nematode-infected plant, i.e. their tolerance rather than resistance increased. The absence of nematodes in the roots of the donor plants rules out any role of nematode-derived compounds in the observed effect. It can only be attributed to the compounds in the leachates derived from the attacked plants. Root exudates are known to carry specific information about biotic/abiotic stress, plant growth and genetic identity (kin recognition) of the donor plants (Bezemer and Van Dam, 2005; Falik et al., 2014, 2012; Semchenko et al., 2014). These studies reported responses in exudate-exposed plants such as altered resistance to aboveground herbivory, stomatal aperture, flowering timing and root morphology change. It is possible that some plant-defensive compounds induced by nematode infection were excreted into the rhizosphere of the nematode-free roots of the same plant and regarded as a reliable cue to induce a tolerance response of overcompensation in the leachate-receiving plants. This is, to our knowledge, the first evidence of a tolerance response of overcompensation induced not in plant attacked by herbivores, but in plant receiving a signal from roots of a herbivore-damaged plant. Why this response was not seen in plants receiving leachates from directly attacked roots is unclear. We speculate that some nematode-derived compounds may cause the plant to abandon this tolerance response. If the attack by nematodes is imminent, as indicated by presences of nematodes in the immediate environment, there may not be enough time for the plant to increase its growth, hence rendering the tolerance response maladaptive.

Plants receiving leachates from locally nematode-infected root systems had a size and belowground root traits similar to those of the plants receiving control leachates. However, the plants receiving leachates from nematode-free and from infected roots of an infected donor plant both developed significantly more shoot tillers than these control receiver plants. This indicates a defensive response in aboveground traits. *Agrostis stolonifera* often reproduces primarily vegetatively, spreading via stolons developed from shoots, which may become separated and continue growing from the stolon nodes as separate plants (Widén 1971). Producing more aboveground shoot tillers may represent an avoidance mechanism for plants by enabling future recruitment of stolons from vegetative shoots to grow away from the nematode-infected area. The nematode-infected plants had twice the amount of nutrients and access to double the amount of soil compared with the control plants in this study. This may have caused an effect on their exudates and ultimately affected the leachates. Splitting the roots may also affect the growth of the nematode-infected plant. Thus, it would be more ideal to also use the split-root system in the control plants to better compare the effect of leachates on the receiver plants with that of leachates from the nematode-infected plant.

Our findings suggest that signals from roots can mediate plant-plant interactions upon root herbivory and elicit different defensive responses in neighbouring plants to adjacent roots with a different locality of root feeders. However, regardless of these differences, neighboring plants share a similar aboveground dynamic of increased tiller production in response to belowground potential threats. We used leachates that mainly differed in root or nematode stylet exudates in our study, as in the field such exudates can also be diffused away from the roots via soil water. This aqueous spread of signals for plant-plant communication in the soil may enable plants to efficiently exchange information on possible belowground stress factors and thus for signal receivers to respond quickly. We did not quantify the chemical composition of these leachates and further studies should focus on compositional differences of root exudates between local nematode-infected roots and distal healthy roots from the same nematode-infected plant. The active distance and time for this aqueous way of communication in the soil should be also considered in future work on this topic.

## Supporting information

Data for the experiment

## Acknowledgements

Financial support from CSC (China Scholarship Council) and the Terrestrial Ecology Unit of Ghent University is gratefully acknowledged. Peihua Zhang is a doctoral fellow supported by CSC (#201506190135).

